# In HSV-1, the LAT Enhancer Drives Pre-IE VP16 Transcription to Initiate Reactivation

**DOI:** 10.64898/2026.01.06.697999

**Authors:** Ziyun A. Ye, Mason A. Shipley, Pankaj Singh, Donna M. Neumann

## Abstract

The HSV-1 VP16 protein, delivered to the nuclei of epithelial cells from the tegument layer of infecting virions, drives viral Immediate-Early (IE) transcription to initiate productive (lytic) infection. Latent HSV-1, established in sensory neurons, reactivates to a lytic infection in the absence of a recent *de novo* infection event that would deliver tegument proteins like VP16. Therefore, the role of the protein VP16 in the exit from HSV-1 latency has been hotly debated for decades. Here we show that VP16 transcription during latency is silenced by CTCF and cohesin proteins bound to a newly identified CTCF insulator site. Upon a reactivation stimulus, CTCF and cohesins are evicted and VP16 is transcribed prior to, and in the absence of, any other viral transcription. We previously identified long-range *cis* spatial interactions between the LAT and VP16 loci of the latent HSV-1 genome. Here, our data further indicate that the LAT enhancer serves as a neuron-specific enhancer of VP16 transcription during reactivation. Collectively, we show the HSV-1 latent genome is hard-wired for reactivation by a long-range spatial interaction between the viral gene encoding the protein that initiates lytic phase transcription (VP16) and the viral enhancer element that drives its transcription (LAT enhancer).

**Author Summary:** Herpes Simplex Virus 1 (HSV-1) is a significant lifelong human pathogen that infects 70% of adults worldwide. HSV-1 establishes a persistent latent infection in the peripheral nervous system where the virus can periodically reactivate in response to environmental stressors. These reactivation events result in recurrence of corneal infections that can lead to corneal blindness. It is becoming clear that CTCF insulators play a key role in the maintenance of gene silencing in DNA viruses that can establish latency. CTCF insulators are essential regulators of chromatin structure and play vital regulatory roles in transcriptional control of DNA viruses by organizing chromatin architecture during both latent and lytic stages of virus lifecycles. Here we have identified and functionally characterized a novel CTCF insulator, located at the transactivating gene VP16. We provide evidence that this insulator is involved in silencing VP16 during latent infection, while eviction of insulator protein coincides with increased VP16 gene expression during reactivation. To our knowledge, these are the first data that show that CTCF insulators is a key regulatory element in the exit from latency in a VP16-dependent manner and has important implications in understanding the heterogeneity of HSV-1 reactivation in different populations of neurons.

## Introduction

Herpes Simplex Virus 1 (HSV-1) is a lifelong human pathogen that infects >70% of the population by early adulthood (1, 2). During the primary infection, HSV-1 infects epithelial cells and spreads to sensory neurons that are innervating the face or eyes and the virus travels by retrograde transport to neuronal cell bodies in the peripheral nervous system to establish life-long latency. Latent HSV-1 is dormant, yet it still presents a significant clinical challenge since latent virus can reactivate in response to stressors (3, 4). During reactivation, virions travel back to peripheral sites, including the cornea, to replicate and cause pathogenesis. Importantly, repeated reactivation in the eye can result in vision loss (1, 5, 6).

Both the primary and reactivation phases of the HSV-1 life cycle result in replication and the production of infectious virions. However, the molecular mechanisms that drive replication during the primary infection in epithelial cells are different from mechanisms driving the initiation and progression of reactivation in neurons. For example, during the primary infection, tegument-delivered VP16 colocalizes in the cytoplasm with the cellular protein Host Cell Factor 1 (HCF-1), where it is then transported to the nucleus of the infected cell. In the nucleus, the HCF-1-VP16 complex binds to the cellular protein Octamer binding transcription factor 1 (Oct-1), and this complex initiates Immediate Early (IE) gene expression, driving the expression of early (E) and late (L) genes to produce progeny virus (7–10). During the primary infection, efficient viral gene expression also requires that histone demethylases remove repressive chromatin modifications that are initially deposited on HSV-1 genomes (7, 9–13).

In contrast, there is no tegument-delivered VP16 to the soma of sensory neurons as the virus establishes latency. Here, epigenetic silencing of viral lytic genes is achieved through host-mediated deposition of heterochromatin marks H3K9me2/3 and H3K27me3 that are maintained until stress response pathways are activated and reactivation progresses (14–19). Reactivation of HSV-1 is a controlled and heterogenous process where only a small number of neurons reactivate at a given time to produce progeny virus and the reversal of latency is likely epigenetic but the mechanism(s) for how HSV-1 achieves this remain an area of scientific debate (3, 4, 20–31). For example, in 2012 Kim, *et. al* reported that following reactivation stressors to quiescently infected primary superior cervical neuronal cultures, HSV-1 reactivation proceeds in a biphasic manner (Phase I/II), where the initial bursts of all 3 kinetic classes of genes are simultaneously observed by 18 hours post-reactivation (hpr) (Phase I) and is independent of VP16 expression and synthesis during Phase I (21). The Cliffe lab later showed in a series of reports that Phase I gene expression was dual leucine zipper kinase (DLK)-dependent and was preceded by a methyl/phospho switch at heterochromatic marks on latent viral genomes to allow for de-repression and subsequent transcription of IE genes in phase I reactivation (3, 32). In contrast, using an *in vivo* model of HSV-1 latency and induced reactivation, Sawtell and Thompson proposed and provided compelling evidence that VP16 is a pre-IE gene in the context of neurons, and the de-repression of the VP16 promoter drives *de novo* synthesis of VP16 and the exit from latency (20, 33, 34). Consequently, the role of the protein VP16 in the exit from HSV-1 latency has been hotly debated for decades.

To further explore the role of VP16 in reactivation and how silencing of VP16 expression is reversed, we utilized the Lund Human Mesencephalic Neuronal cell line (LUHMES)—an embryonic neuronal precursor cell line that can be differentiated to postmitotic neurons that mimic key aspects of HSV-1 latency and reactivation (35)—to determine the kinetics of VP16 expression following the initiation of reactivation. LUHMES were infected HSV-1 and after establishing a quiescent infection were reactivated with the PI3 kinase inhibitor wortmannin. Following the initiation of reactivation, VP16 expression significantly and rapidly increased even prior to IE gene expression. Moreover, this increase in gene expression was unaffected by the presence of the protein synthesis inhibitor cyclohexamide, indicating VP16 transcription was independent of *de novo* viral protein synthesis. Companion animal experiments using mice and explant induced reactivation were consistent with the LUHMES model for VP16 expression, confirming that our findings were not LUHMES specific. While it was clear that VP16 gene expression rapidly increased in both models, mechanistic insight into how the reversal of VP16 silencing or de-repression of the VP16 promoter occurred remained elusive.

Cellular genomes are organized into transcriptionally active and repressive chromatin domains and insulators separate and maintain these epigenetic domains (36). Chromatin insulators bind the CCCTC binding factor, or CTCF, a ubiquitously expressed zinc finger DNA-binding protein that binds to specific sequence motifs (37). In mammalian cells, CTCF insulators are the master regulators of transcription and function as enhancer-blockers or as barrier elements (36–38). CTCF insulators also control transcription three-dimensionally (3D) by forming chromatin loops that coordinate long-range spatial interactions between distal promoters and enhancers for transcriptional control (37, 39–41). Like cellular genomes, latent HSV-1 genomes are also organized into transcriptionally active and repressive domains that are flanked by CTCF insulators, suggesting that CTCF insulators maintain gene silencing during latency (42–44). Early computational analysis identified seven putative CTCF binding sites in HSV-1 genomes, six of which were in the repeat regions encompassing the immediate early genes ICP0 and ICP4, as well as LAT 5’exon region of the HSV-1 genome while the seventh was in the unique short region (42). The locations of each of these elements appears deliberate for function, as they all flank reactivation-required genes including the LAT enhancer element (LTE) (45, 46). These binding sites were subsequently shown to be enhancer-blocking or barrier insulators that were enriched in CTCF proteins during latency (24, 44). We later showed that siRNA-targeted CTCF depletion in latently infected rabbit neurons led to spontaneous reactivation, supporting that CTCF proteins must be bound to the viral genome to maintain latency (27). In contrast, CTCF protein was differentially evicted from virally encoded insulators at the earliest time measured (1-2 hpr), suggesting that CTCF insulator function must also be disrupted for the progression of reactivation (23, 24). Consequently, CTCF eviction precedes the accumulation of lytic transcripts described in the biphasic reactivation model (21) and coincides with increased VP16 gene expression in the data presented here, suggesting that CTCF eviction may also be a pre-requisite of the initial lytic gene expression cascade observed during reactivation.

We recently used Circular Chromosome Conformation Capture-Sequencing combined with Unique Molecular Identifiers (UMI)-4C to quantitate multiple long-range *cis-*interactions in quiescent HSV-1 genomes, including one interaction that mapped to sites near the VP16 and LAT loci (47). This finding suggested that this long-range spatial interaction likely spatially positioned the VP16 promoter to the distal LTE possibly for transcriptional control. Considering that derepression/ activation of the VP16 promoter might be dependent on the LTE, we used a recombinant virus with a 307 base pair deletion of the LTE (17ΔA) to determine if the LTE meaningfully contributed to VP16 expression. We found in both quiescently infected LUHMES and *in vivo* that in the absence of the LTE, there is no increase in VP16 gene expression following reactivation. The absence of the LTE also impacted the expression of genes from other kinetic classes but it was unclear whether this was due to VP16 expression or a consequence of the absence of the LTE. To resolve this, we generated reporter constructs and showed that the LTE was required for VP16 promoter activation in neuronal cells but had no effect on other viral gene promoters tested, supporting that VP16 expression following the initiation of reactivation is LTE-dependent.

The identification of long-range spatial interactions between VP16 and LAT loci suggested that there was an unidentified CTCF insulator near VP16 that was nucleating this long-range 3D interaction. Therefore, we performed computational analysis to identify putative insulators near the VP16 locus and then generated a series of reporter constructs to test the enhancer-blocking activity of each of the putative insulator sequences to the LTE (48, 49). We identified one binding motif near VP16 (subsequently named CTUL1) as an enhancer blocker. Conventional ChIP-qPCR quantitated significant enrichment of both CTCF and cohesin proteins to this unique site, further supporting its role as a functional CTCF insulator that forms long-range spatial interactions to maintain latency. Following induced reactivation, we found that CTUL1 completely lost enrichment of CTCF and cohesin by 2 hpr (the earliest time measured by ChIP) and the loss of CTCF coincided with increased IE, E and L gene expression in both LUHMES and mice trigeminal ganglia (TG). We then tested whether the CTUL1 insulator could block VP16 promoter activity and showed that the CTUL1 insulator blocked LTE driven VP16 promoter expression neuronal cells and our data suggest that the LTE serves as a neuron-specific enhancer of VP16 transcription during reactivation. Collectively, we show the HSV-1 latent genome is hard-wired for reactivation by a long-range spatial interaction between the viral gene encoding the protein that initiates lytic phase transcription (VP16) and the viral enhancer element that drives its transcription (LTE).

## Results

### VP16 expression is detected “pre-IE” following the initiation of reactivation

Sawtell and Thompson previously reported that *de novo* synthesis of VP16 initiated the exit from latency in mice sensory neurons (33, 34). To test this in the context of gene expression, we quiescently infected LUHMES cells with 17*Syn*+ (Genbank NC_001806) and then induced reactivation using the PI3 kinase inhibitor wortmannin (35). Total RNA was collected from LUHMES cells at 0, 1, 3, 6, and 24 hpr and qRT-PCR was used to quantitate VP16 expression. Viral RNA transcripts of representative genes from all kinetic classes were normalized to host GAPDH and are displayed as a fold change relative to latency, which is set to 1. Here, it is also important to note that baseline levels of expression of all lytic genes are exceptionally low during latency with C(t) values in qRT-PCR falling above the threshold for what is considered a negative value (C(t) above 36). These relative values are used for the quantitation of lytic genes for comparison purposes only. In our PCR experiments, VP16 expression significantly increased compared to baseline latent levels at the earliest time measured following the initiation of reactivation (1 hpr-**Fig. 1A**). The RNA transcript level further increased by 3 hpr and remained significantly higher than latency through 24 hpr, consistent with reactivation. In contrast, the IE gene ICP27 and E gene ICP8 did not have significant increases in expression until the 3 hpr timepoint (**Fig. 1A**), and the late genes VP5 and gC were not expressed until 24 hpr, as expected during a reactivation event (**Fig. 1A**). Classified as a late gene in the primary infection, here VP16 expression increased much earlier than VP5 and gC from the same kinetic class, and earlier than the IE gene ICP27 upon reactivation. This was an exciting finding suggesting that in the context of reactivating genomes in neuronal cells VP16 expression is “pre-IE”. These findings in LUHMES cells were consistent with increased viral gene expression quantitated following explant induced reactivation in latently infected mice TG (**S Fig. 1**).

**Figure 1:**
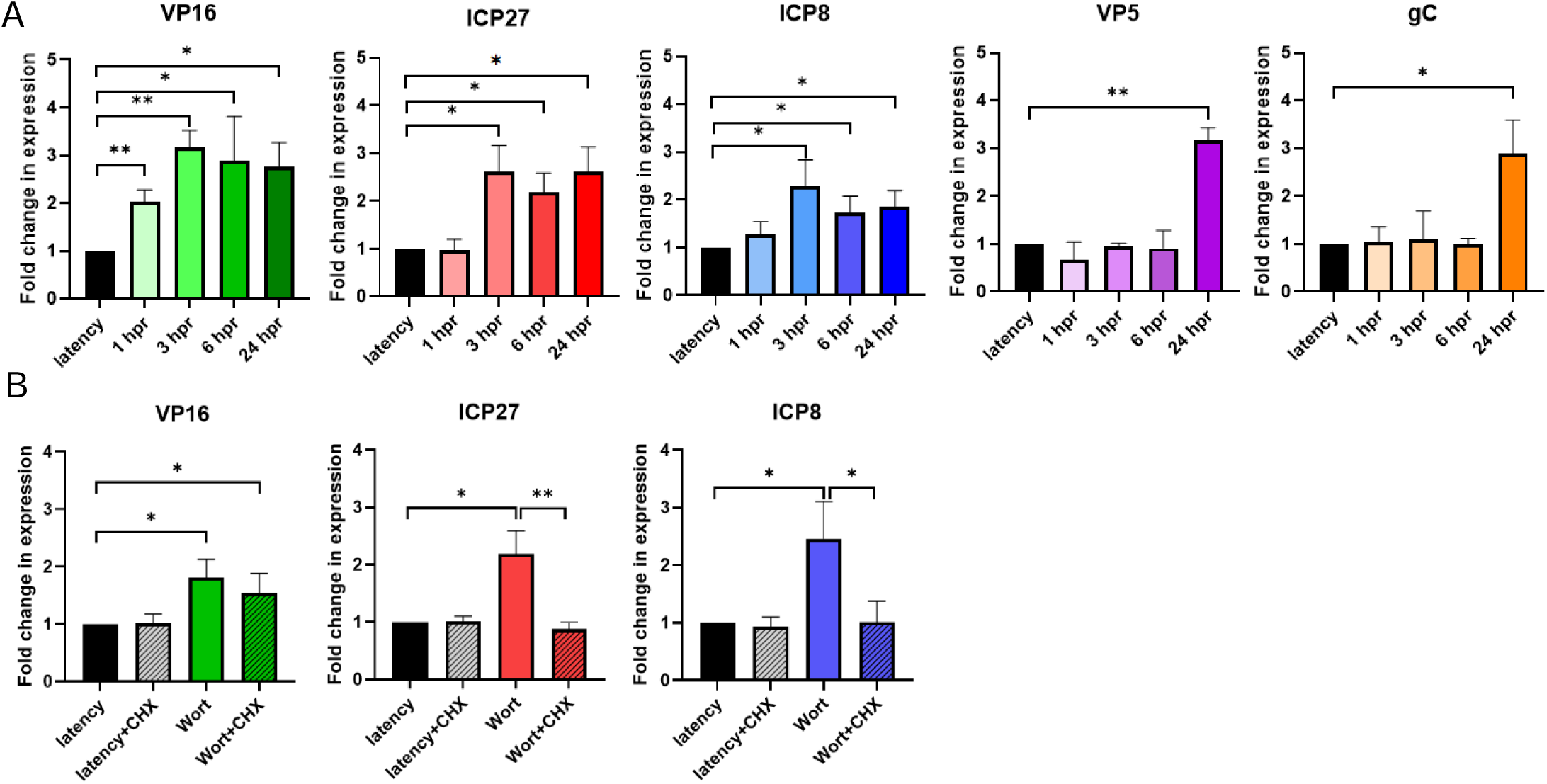
Expression of VP16 and representative genes from each kinetic class were quantified by RT-qPCR in LUHMES cells. **A**. Cells were quiescently infected at an MOI of 0.2, and total RNA was extracted at 0, 1, 3, 6, and 24 hours post Wortmannin-induced reactivation. Gene expression was quantified using primers and probes specific to each gene tested (see Table S2). Relative expression values for each viral gene were normalized to host GAPDH expression and were plotted as fold change in expression relative to latency (set to 1). n=3. \**p*<0.05, \*\**p*<0.005, \*\*\**p*<0.0005 following Student’s t-tests. **B**. Cells were quiescently infected at an MOI of 0.2. Total RNA was extracted at 0 and 6 hours post Wortmannin-induced reactivation in the presence and absence of cycloheximide (CHX) treatment. Relative expression values for each viral gene were normalized to host GAPDH expression and were plotted as fold change in expression relative to latency (set to 1). n=3.

During the primary infection, tegument-delivered VP16 transactivates the expression of IE genes in the kinetic cascade of gene expression. Our data here suggest that VP16 expression increases prior to IE genes during reactivation, suggesting that downstream VP16 protein synthesis is also prerequisite for the initiation of IE gene expression during reactivation episodes. To test this, we inhibited viral protein synthesis using cycloheximide (CHX) and quantitated lytic gene expression by qRT-PCR following reactivation. Reactivation was induced by wortmannin and transcript levels were quantitated at 6 hpr in the presence and absence of CHX. CHX treatment did not alter baseline levels of viral transcripts in latency (**Fig. 1B**). At 6 hpr, VP16, ICP27 and ICP8 had significant transcription induced by wortmannin alone, consistent with our earlier findings. In contrast, transcription of ICP27 and ICP8 were blocked in the presence of CHX (**Fig. 1B**), indicating that IE and E gene expression in reactivation requires *de novo* protein synthesis first. Notably, CHX had no significant effect on VP16 transcription following reactivation. This result suggests that VP16 transcription is independent of *de novo* viral protein synthesis and likely be upstream of the transcription cascade of IE, E and L genes in reactivation. We further validated VP16 protein was detectable at 6 hpr in the LUHMES infection model by immunofluorescence microscopy (**S Fig. 2B**). Here, VP16 protein can be seen localized in the nuclei of ∼15-20% of cells induced to reactivate, further indicating that *de novo* synthesis of VP16 is an early and fundamental step in reactivation of HSV-1.

### VP16 fails to express at early times following reactivation in the absence of the LAT enhancer (LTE)

LTE, the enhancer element encoded in the 5’ exon of LAT, is required for efficient HSV-1 reactivation (18, 45), albeit through unidentified mechanisms. Our previously published work identified a long-range spatial interaction in quiescent HSV-1 genomes mapping to the LAT and VP16 loci, and we hypothesized that this 3D organization was maintained for transcriptional purposes during reactivation. To test this further, we utilized a recombinant HSV-1 strain, generated from the 17*Syn*+ background containing a 307 bp deletion of the LTE (n.t.119,200-119,507) (17ΔA- a kind gift from the Bloom lab- UFL). The 17ΔA recombinant was sequenced prior to use and contained no secondary mutations. We routinely use wt parental viruses for our controls because, while rescue viruses are useful in proving that the genetic lesion is responsible for a change in phenotype, the process of generating the rescue results in genetic drift of the rescue from the wt virus following the transfection and multiple rounds of plaque purification. It is possible that these genetic drifts can result in significant differences in the 3D genome structures or chromatin architecture between the wt and rescue viruses. For these reasons, we propose that recombinant virus comparisons to wt virus are the more robust control for our experiments. Further, we phenotyped the recombinant prior to use in Vero cells and determined there were no growth or replication defects in the lytic infection (**S Fig. 3A**). In LUHMES cells and *in vivo*, 17ΔA established latency equivalent to the parent wt strain 17*Syn*+ both *in vitro* and *in vivo* (**Fig. 2A; S Fig. 3B**). To compare gene expression during reactivation, LUHMES were infected with either 17ΔA or wt virus. Following the establishment of latency, cells were subjected to wortmannin-induced reactivation at 1, 3, 6, and 24 hpr. qRT-PCR showed that in the absence of the LTE, there was no increase in VP16 transcription in the absence of the LTE until 24 hpr (**Fig. 2B**- consistent with the *in vivo* data shown in **S Fig. 3)** and even then, this increase in transcription did not correspond to true reactivation where the production of infectious virions could be detected (**S Fig. 4**). This finding and its implications are further discussed in the Discussion portion of this manuscript. Further, and consistent with our findings that VP16 expression is a prerequisite for IE gene expression, we could no longer detect increased transcription of ICP27 in the absence of the LTE (**Fig. 2C**). Finally, increased viral transcription in representative E and L gene classes was absent in the recombinant virus and gC had a significantly lower expression level than the latency baseline, implying transcript degradation was occurring over time. These findings, combined with the inability to reactivate in the absence of the LTE confirm that these are not delayed expression kinetics, rather that VP16 expression is a first step in reactivation and is driven in a LTE-dependent manner.

**Figure 2:**
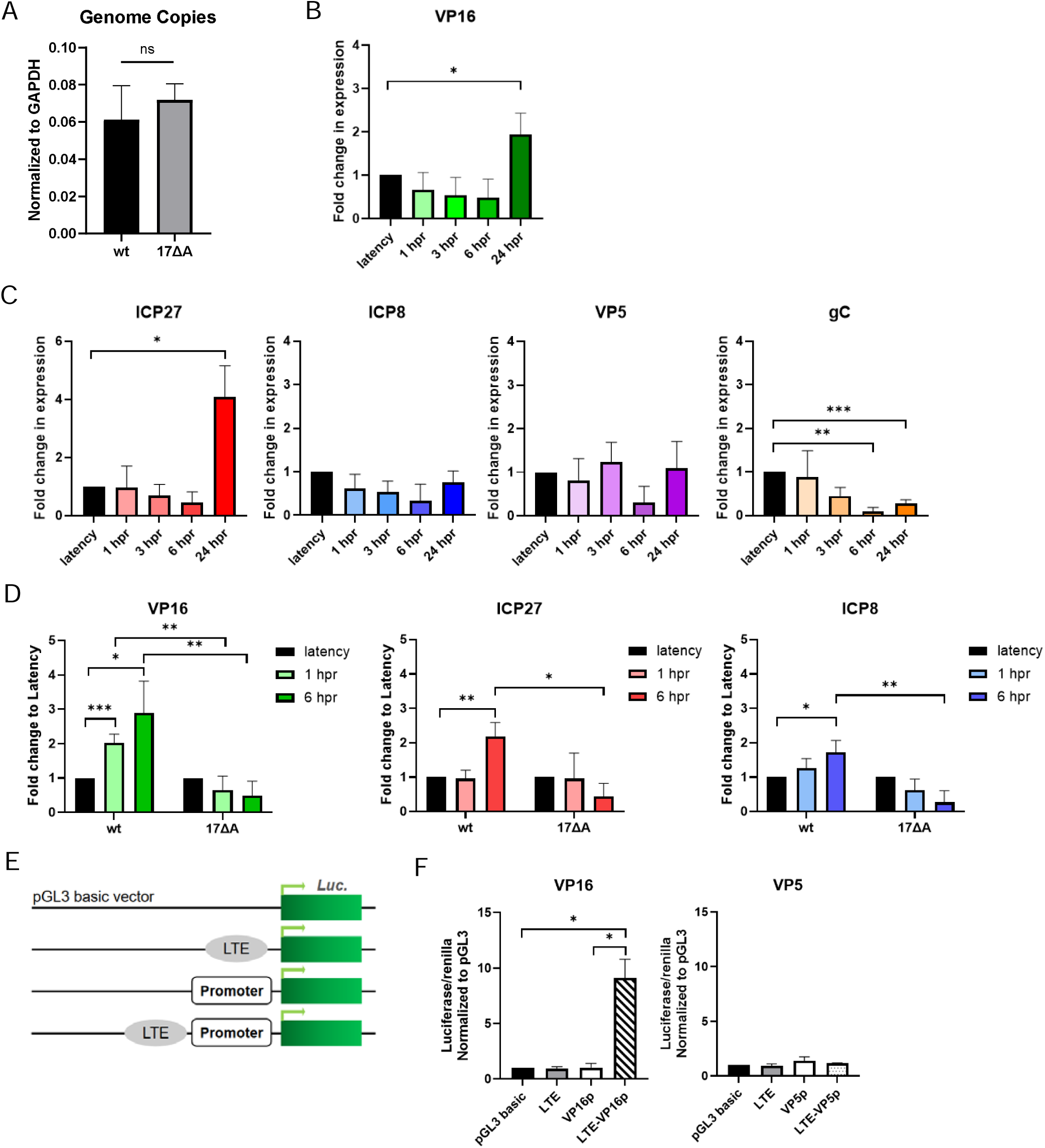
17ΔA gene expression following the application of reactivation stimulus wortmannin. **A.** wt and 17ΔA HSV-1 genome copies in quiescently infected LUHMES cells (MOI of 0.1). Relative genome load was determined by qPCR copy number of DNA polymerase normalized to that of host GAPDH. (n=3). Statistical significance between wt and 17ΔA was calculated by unpaired two-tailed Student’s t-tests with equal variance. **B.** VP16 gene expression in 17ΔA was quantified by qRT-PCR in LUHMES cells. Cells were quiescently infected at an MOI of 0.2 and total RNA was extracted at 8 dpi (latency), 1, 3, 6 and 24 hpr. Gene expression was quantified using primers and probes listed in Table S2. Relative values for each gene were normalized to host GAPDH expression and were plotted as fold change in expression relative to latency (set to 1). n=3. \**p*<0.05, \*\**p*<0.005, \*\*\**p*<0.0005 following Student’s t-tests. **C**. Expression of representative genes from each kinetic class were quantified by RT-qPCR as described in panel B. **D**. Comparison between gene expression of wt and 17ΔA at 1 and 6 hpr. **E.** Luciferase reporter constructs were generated to test the activation of VP16 promoter in neurons using the promoterless pGL3 basic vector, pGL3 basic-LTE (n.t.118,889-119,478), VP16 promoter only, VP5 promoter only, and VP16 or VP5 promoter inserted into the LTE plasmid. **F.** Luciferase reporter assays were performed in mice N2a cells. All transfections were completed in triplicate wells and were repeated three times biologically. All luciferase values were normalized to the pGL3 basic vector (set to 1). Statistics were determined by unpaired two-tailed Student’s t-tests with unequal variance.

*In vivo*, we infected mice with either wt or 17ΔA HSV-1 and established latent infections. Mice TG were harvested at 28 dpi (latency) and explant was performed at 2, 6, and 48 hpr. After total RNA extraction, representative genes were analyzed by qRT-PCR for expression levels. Gene expression in both wt and 17ΔA were consistent with our LUHMES data, namely that early expression of VP16 and ICP8 in reactivation was suppressed in 17ΔA (**S Fig. 3C**). To compare between the two viruses, fold change expression of the wt and recombinant viruses were compared at 1 and 6 hpr and combined on one graph (**Fig. 2D**). To quantify replication in both viruses, we extracted DNA from quiescently infected LUHMES at latency and 48 hpr. We observed no increase in genomes in the 17ΔA infected cells compared to latency, consistent with attenuated reactivation (**S Fig. 4A**). Plaque assays done on the supernatant of LUHMES infected with the mutant virus and subjected to reactivation showed the absence of progeny virus (**S Fig. 4B**), supporting that the deletion of the LTE inhibits reactivation altogether.

### The LTE drives VP16 promoter activity in neuronal cells

We detected increased transcription of VP16 at 1 hpr. Nonetheless, little is known about mechanisms that initiate expression of VP16 in neurons in the context of HSV-1 reactivation. Based on our previous work (47), we hypothesized that the VP16 promoter is activated in an LTE-dependent manner sequence in neuronal cells. To further test this, luciferase reporter constructs were generated by inserting either the VP16 promoter or the VP5 promoter, another late gene, into the promoterless pGL3-basic vector, in the presence or absence of the LTE (**Fig. 2E**). Reporter constructs were transfected into mice Neuro-2a (N2a) cells. Following transfection, luciferase expression was quantitated for each of the reporter constructs as described in the Methods section. Introduction of the LTE sequence alone did not promote luciferase expression. Interestingly, neither the VP5 nor the VP16 promoter constructs resulted in luciferase expression in N2a cells (**Fig. 2E**). However, the expression from VP16 promoter was enhanced nearly 10-fold in the presence of the LTE, indicating that the VP16 promoter requires the LTE for activation in the context of neurons. By comparison, luciferase expression from the VP5 promoter was still absent in the presence of LTE, suggesting that the LTE is not a promiscuous enhancer for just any HSV-1 promoter. In contrast, in HEK293FT cells the VP16 promoter alone significantly increased luciferase expression, and the addition of the LTE enhanced this expression >2-fold, further supporting that the LTE has cell-type specific regulatory functions toward the VP16 promoter (**S Fig. 5**).

### A novel enhancer-blocker identified in the UL region of HSV-1 is located downstream the VP16 promoter

Early sequence algorithms targeting the identification of CTCF insulators in HSV-1 scanned the genome for reiterated CCCTC/CTCCC motifs. None were identified near the VP16 locus by these early analyses. However, CTCF proteins can bind to motifs other than CCCTC/CTCCC and considering the importance of CTCF insulators to transcriptional control, we anticipated that there were likely unidentified DNA binding sequences that bound the CTCF protein near VP16. To test this, we performed a comprehensive search of both upstream and downstream of the VP16 locus for potential binding motifs using an online CTCF binding site prediction tool. The CTCFBSDB 2.0 program is a comprehensive collection of experimentally determined and computationally predicted CTCF binding sites (48, 49). Following the input of a genomic sequence, the software generates predicted motif locations, site orientation (direct or indirect strand of DNA) and a compatibility score within that genomic sequence. Compatibility scores greater than 3.0 are considered the most highly probable binding motifs for CTCF. Using a ∼2 kb input sequence of the HSV-1 genome (n.t. 101,200-103,600 from strain 17*Syn*+) we identified four individual (non-overlapping) putative CTCF binding sequences, labelled A-D (**Fig. 3A, Table S1**). Reporter constructs were made using the pGL3-control vector (Promega) backbone that contains an SV40 promoter-driven luciferase gene (**Fig. 3B**). First, we generated a plasmid with a 580 base pair insertion that corresponded to the LTE of HSV-1 (n.t. 118,888-119,477), as previously described (24). The A-D reporter constructs were generated by inserting the A, B, C or D sequences identified by the CTCFBSDB program between the LTE and SV40 promoter in the LTE plasmid (**Fig. 3B**). Each reporter construct contained predicted CTCF binding sequence and an additional 150-350 bp of nucleotides upstream and downstream of the predicted binding motif. Transient transfections of each individual reporter construct were done in mice N2a cells to determine enhancer-blocking activity. Following transfection, luciferase expression was quantitated for each of the reporter constructs as described (24). Controls for each expression assays were the pGL3-control vector (negative control) and the previously published CTRL2 plasmid (positive control) (24, 42). Consistent with previously reported data, there was a 4-fold increase in luciferase expression in the LTE containing plasmid (**Fig. 3C**) (23, 24, 42). We then quantitated luciferase expression for the reporter constructs containing the LTE and sequences A-D and found that only one of the constructs significantly decreased expression compared to the LTE reporter, suggesting this element acted as an enhancer-blocker to the LTE in neuronal cells (D- near the VP16 promoter; **Fig. 3C**). In contrast, there was no observed inhibition of the LTE to driving luciferase expression in any of the other plasmids that contained inserts of the same size and GC content, confirming that the activity of the D insert was not due to the distance of separation between the LTE and SV40 promoter in the pGL3 vector.

**Figure 3.**
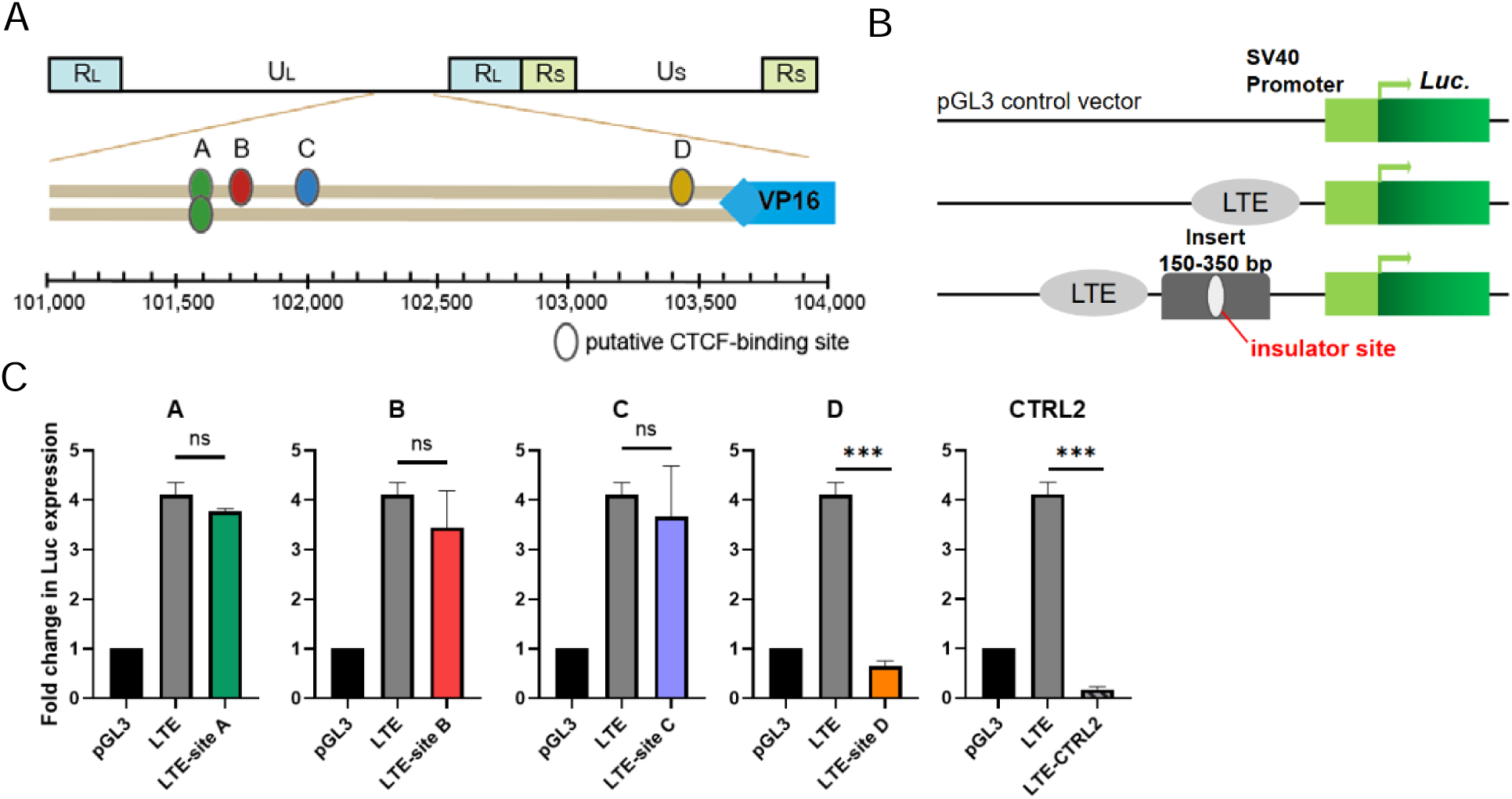
Luciferase reporter constructs were generated to test the enhancer-blocking ability of the predicted CTCF insulator sites. **A**. Schematic representation of CTCFBSDB predicted CTCF binding sites in the UL region of the HSV-1 genome near VP16. **B**. Representative orientations for each of the constructs tested include the commercially available pGL3-control vector with an SV40 promoter and a luciferase gene as the plasmid backbone; the LTE inserted into the control vector; each putative CTCF-binding site identified in Table S1 together with ∼150-350 bp flanking sequences inserted downstream of the LTE. **C**. Luciferase reporter assays were performed in N2a as a representative neuronal cell line. All transfections were completed in triplicate. Data represent at least three biological replicates. All luciferase values were normalized to the pGL3-control vector as a fold change in expression relative to control (set to 1). CTRL2 served as a positive control with known LTE-blocking activity in N2a. Statistical comparisons were done on fold changes between the LTE construct and the LTE-CTCF site. \**p*<0.05, \*\**p*<0.005, \*\*\**p*<0.0005 by unpaired two-tailed Student s t-tests with unequal variance.

### CTCF and cohesin proteins are enriched on the CTUL1 site during latency

To maintain consistency with the insulator nomenclature that was previously published (42) the sequence identified as an enhancer blocker (site D) was named **CTUL1.** Since functional insulators bind the CTCF protein, we determine if CTCF protein was bound to **CTUL1** in the LUHMES cell culture model of HSV-1 quiescence. Chromatin Immunoprecipitation (ChIP) assays combined with qPCR was done using primers and probes adjacent to nucleotide sequences of the identified CTCF binding sites (**Table S2**), as previously described (23, 24, 50). Prior to analyses, each of the ChIP experiments were validated using the positive control host H19/Ifg2 and only samples with >5-fold enrichment of CTCF at the host locus versus IgG were further analyzed (**Fig. 4A**). As a positive viral control for CTCF binding on the HSV-1 genome, we selected the CTRL2 insulator, a functional insulator located downstream from the LTE that is significantly enriched in CTCF during latency (23, 24). Quantitation of CTCF enrichment at the newly identified CTUL1 site showed the motif was significantly enriched in CTCF protein relative to IgG (>10-fold enrichment- **Fig. 4A**). On the other hand, we detected no significant CTCF enrichment at the HSV-1 glycoprotein C locus (gC), a site with no predicted or known CTCF binding domains, which served as a negative control.

**Figure 4:**
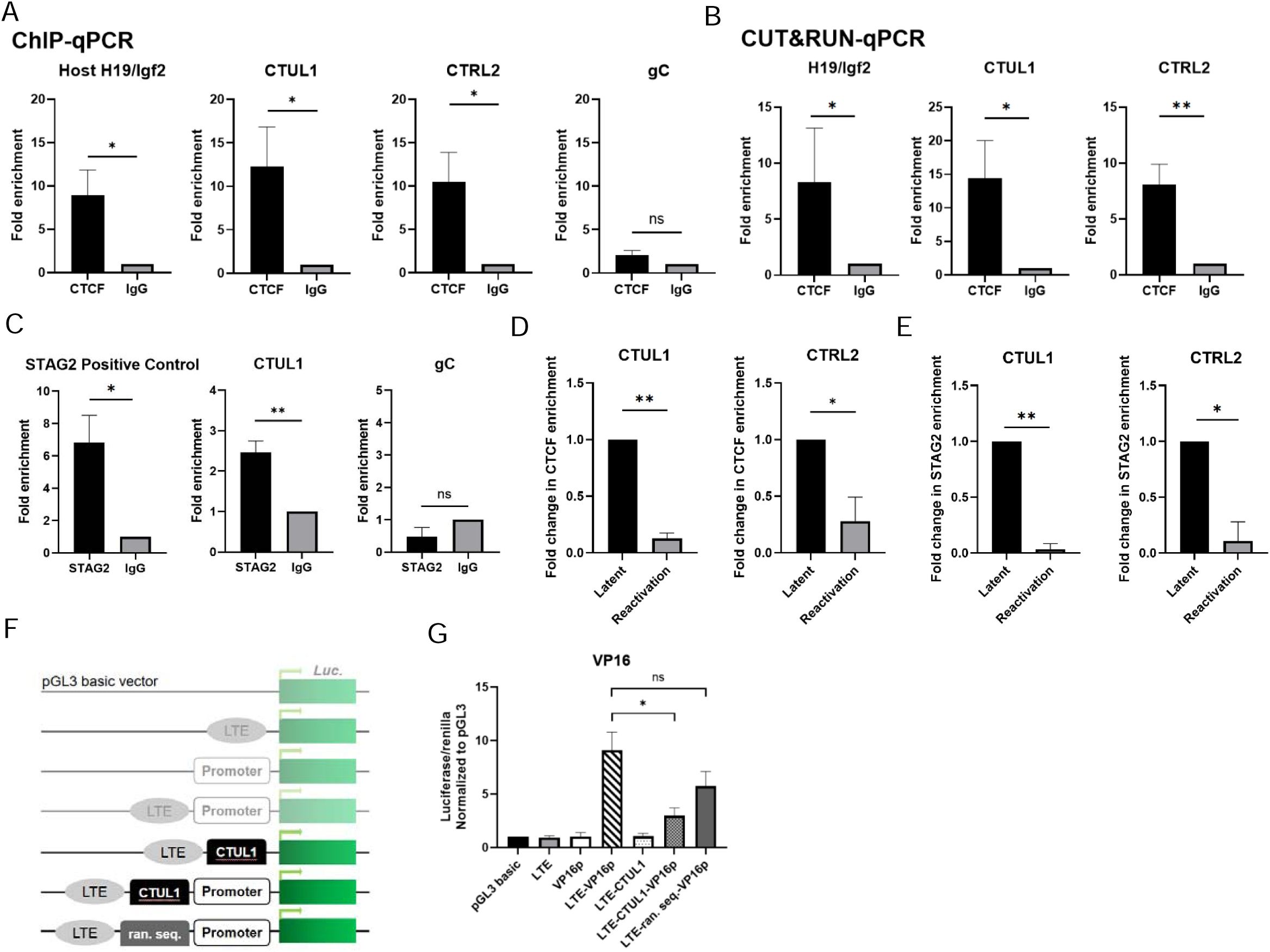
CTCF and cohesin proteins are enriched on CTUL1 during HSV-1 latency but evicted by 2 hpr. Protein binding was determined by ChIP or CUT&RUN and quantified by qPCR using primers and probes specific to the given gene regions of HSV-1 (Table S2). All relative values for CTCF enrichment were normalized to those of the non-specific binding control IgG and were plotted as fold enrichment relative to IgG (set to 1). n=3-5. **A.** LUHMES cells were infected at MOI of 0.3 and harvested at 8 dpi for ChIP. CTCF binding in LUHMES was validated at the host positive control region H19/Igf2. qPCR primers specific to CTUL1 or CTRL2 were used to quantitate CTCF enrichment (Table S2). gC served as a negative control of CTCF binding on HSV-1 genome. Each ChIP assay represented one 6-well plate of cells. n=3. Binding significance was determined by Student’s t-tests. \**p*<0.05. **B.** CUT&RUN-qPCR was performed with quiescently infected LUHMES cells as described above. CTCF binding significance was determined by one-way analysis of variance (ANOVA). n=3. **C.** ChIP-qPCR was done with the anti-STAG2 antibody on latently infected LUHMES. STAG2 binding in LUHMES was validated at a host positive control region on human Chromosome 1, as previously reported (Table S2). Primers specific for CTUL1 were used to quantitate STAG2 on each site following ChIP. n=4. gC served as a negative control on the viral genome without STAG2 enrichment. Binding significance was determined by unpaired two-tailed Student’s t-tests. \**p*<0.05, ** *p*<0.005. **D**. LUHMES cells were infected at MOI of 0.3 for 8 days and subjected to wortmannin treatment for 2 hours, followed by ChIP-qPCR. CTCF bound/input values at CTUL1 and CTRL2 were normalized to bound/input at host H19/Igf2. Then, fold change was calculated between 2 hpr and latency (set to 1). \**p*<0.05, \*\**p*<0.005 by Student’s t-test. **E**. ChIP-qPCR done with anti-STAG2 antibody on LUHMES subjected to wortmannin treatment for 2 hours. Fold change values plotted were calculated by setting latency to 1. n=3. **F**. Luciferase reporter constructs were generated to test CTUL1 as an enhancer-blocker to the VP16 promoter in the presence of the CTUL1 insulator binding site and the LTE. A 294-bp random sequence was inserted in place of CTUL1. **G.** Transient transfections were performed in N2a cells. All transfections were completed in triplicate wells and were repeated three times biologically. All luciferase values were normalized to the pGL3-basic vector (set to 1). * *p<*0.05 by unpaired two-tailed Student’s t-tests with unequal variance.

To add rigor, we adapted the *in-situ* chromatin profile experiment Cleavage Under Targets and Release Using Nuclease (CUT&RUN) combined with qPCR to assay CTCF enrichment during latency in LUHMES cells. CUT&RUN captures DNA-protein interactions in the native local chromatin environment at a higher resolution (∼150 bp) than conventional ChIP assays. Assays were validated using the positive host control H19/Igf2 as previously described (**Fig. 4B**). Following validation of each CUT&RUN assay, qPCR using primers specific to CTUL1 or the CTRL2 binding motifs were used to quantitate CTCF enrichment. Our data using CUT&RUN-qPCR was consistent with our findings utilizing ChIP-qPCR in quiescently infected LUHMES, further confirming that CTUL1 was enriched in CTCF protein during latency. Collectively, these data confirm that CTUL1 is a functional chromatin insulator in the HSV-1 genome during latency.

CTCF insulators, together with the cohesin complex form and anchor long-range genome interactions, also known as chromatin loops. The cohesin complex is made up of four core proteins (SMC1, SMC3, Scc1 (Rad21) and SA (STAG)) that form a ring-shaped structure that colocalizes with HSV-1 CTCF insulators (51). To determine if cohesin proteins could be detected near the newly identified CTUL1, we quiescently infected LUHMES cells and performed ChIP-qPCR, as previously described, using primers that targeted unique sequences near CTUL1 (**Table S2**). ChIP assays were performed using an antibody against the cohesin complex subunit STAG2 (SA2). The positive cellular control the STAG2 antibody was selected from a region on human chromosome 1, as previously described to ensure the pull-downs were successful (**Fig. 4C**) (52). We then quantitated the enrichment of STAG2 at CTUL1 in quiescently infected LUHMES cells and found a significant enrichment in STAG2 at the CTUL1 insulator. No significant STAG2 enrichment was detected at gC, the negative viral control. These data suggest that the CTUL1 insulator is mediating a long-range spatial interaction near the VP16 locus.

### CTCF and cohesin proteins are rapidly evicted at early times following the application of reactivation stimuli on the CTUL1 insulator

We previously showed that CTCF was evicted from the HSV-1 genome at early time post-reactivation in mice latently infected by wt 17*Syn*+ virus but not in mutants with attenuated reactivation phenotypes, suggesting that CTCF eviction is an important part of HSV-1 reactivation (23). Here we showed that CTCF is enriched at CTUL1 during latency, suggesting the insulator function maintains transcriptional silencing of VP16. We hypothesize that loss of insulator function at CTUL1 through the eviction of CTCF may facilitate VP16 expression. To test if CTCF enrichment is lost at this site after reactivation is initiated, we performed ChIP-qPCR in LUHMES cells using wortmannin as a reactivation stimulus (35). Quiescently infected LUHMES cells were incubated in the presence of wortmannin for 2 hours and then cells were harvested and crosslinked for ChIP. Samples were validated with host control H19/Igf2, which had no significant loss in CTCF during reactivation (**S Fig. 6A**). All enrichment values on viral sites were normalized back to the host control for comparisons across conditions. Graphs were plotted as fold change in CTCF enrichment compared to latency, where latency enrichments were set to 1. When we compared the CTCF enrichment at the CTUL1 insulator at 2 hpr, we found a significant >10-fold decrease in enrichment compared to latency (**Fig. 4D**) and we could not detect differences between the CTUL1 site and IgG, suggesting that CTCF was completely evicted from this site. This is consistent with the positive control CTRL2, which lost CTCF enrichment by 2 hpr (23, 24). We also performed ChIP-qPCR with anti-STAG2 in quiescently infected LUHMES treated with wortmannin to determine whether cohesin proteins are lost at the CTUL1 insulator post-reactivation. While there was no loss to STAG2 enrichment at the cellular control (**S Fig. 6B**), we found significant decreased enrichment at both the CTUL1 and CTRL2 insulators (**Fig. 4E**), suggesting that either the long-range interaction is disrupted at these two sites or that alternative long-range interactions are formed possibly through loop extrusion mechanisms during reactivation (53).

### CTUL1 blocks VP16 promoter activation from LTE in neuronal cells

To test the enhancer-blocking activity of CTUL1 in the context of VP16 promoter, we generated additional reporter constructs that placed the CTUL1 binding motif between the LTE and the VP16 promoter (**Fig. 4F**). Using transient transfections in N2a cells we found that the CTUL1 binding domain suppressed VP16 promoter activity (**Fig. 4G**). To make sure the suppression was not due to extended distance between the enhancer and promoter, we inserted a random sequence near CTUL1 (**Table S1**) between LTE and VP16 promoter. No significant suppression of LTE-driven luciferase activity was observed in this LTE-random sequence-VP16p vector, confirming the enhancer-blocking activity of CTUL1 is sequence-, but not distance-, dependent. This result further supports the hypothesis that CTUL1 has the capacity to silence VP16 expression during latency. Altogether, these data support that CTCF insulators located near VP16 could promote gene expression following CTCF eviction in a LTE-dependent manner.

## Discussion

CTCF insulators are essential regulators of chromatin structure and gene expression in mammalian cells (36, 37). It is becoming increasingly clear that these cellular insulator elements also play vital regulatory roles in transcriptional control of herpesviruses as well. In gamma herpesviruses Epstein-Barr Virus (EBV) and Kaposi’s sarcoma-associated herpesvirus (KSHV), and in beta herpesviruses like HCMV, CTCF insulators regulate the expression of viral genes during latency and reactivation by influencing the three-dimensional structure of the viral episome (39, 54–62). CTCF insulators also contribute to the repression of herpesvirus lytic genes by preventing enhancer-promoter interactions that would otherwise activate viral replication. In EBV, for instance, CTCF binding sites are located near important lytic genes, and these sites help to maintain a repressive chromatin environment around these genes. This silencing is crucial for the virus to remain latent within the host cells. Despite advances made in understanding the role of CTCF insulators in beta and gamma herpesvirus gene silencing and activation, their role has been less defined in how they silence HSV-1 or how they contribute to the exit from latency, the progression of reactivation, or both.

The reversal of latency is epigenetic in nature and driven by cellular factors that transiently reverse viral gene silencing (21), albeit through under-defined mechanisms. We have long-since hypothesized that CTCF insulators maintain the integrity of the silenced HSV-1 genome during latency in the neuron and that insulators must be disrupted, either through eviction or rearrangement, for the virus to exit latency and reactivate. Like cellular genomes, latent HSV-1 genomes are organized into transcriptionally active and repressed chromatin domains flanked by CTCF insulators and early computational analysis of latent HSV-1 identified seven putative CTCF binding sites that encompass regions important for reactivation, including ICP0, ICP4, and LTE. We subsequently showed that at least one HSV-1 encoded CTCF insulator controls IE silencing during latency (63). Furthermore, we have shown that CTCF is evicted from HSV-1 very early after the application of reactivation stimuli and that depletion of CTCF in neurons latently infected with HSV-1 results in reactivation (23). Taken together, these factors support a key role for CTCF insulators in latency and reactivation of HSV-1.

The exit from HSV-1 latency is a highly controlled and heterogenous process where only a small number of neurons reactivate at any given time. Primary neuronal culture models that mimic *in vivo* latency have provided molecular details on the process of HSV-1 reactivation, including evidence that viral mRNAs accumulate in a biphasic manner following the pharmacological inhibition of PI3 kinase that drives synchronous HSV-1 reactivation in primary neuronal cultures. In Phase I of the biphasic model, abundant lytic transcripts from all 3 kinetic classes accumulate simultaneously in the absence of viral protein synthesis or reversal of heterochromatic marks on viral genomes by ∼18 hour post-stimulus (3, 64). VP16, the major transactivating protein that drives lytic gene expression, is not detected in the nucleus of neurons until Phase II (∼48 h post-stimulus) where VP16 is then required for viral replication. In Phase II of the biphasic model, DNA amplification and the production of infectious virus occurs (21). While these reactivation events are well-described in the literature, robust accumulation of lytic transcripts is not apparent until after 18 hpr and does not account for other rigorous scientific data reported in the literature, including the observation that increased IE gene expression is detected by 4 hpr (23, 65), that changes to chromatin around the IE genes can be quantitated by 4 hpr (65) and that *de novo* synthesis of VP16 is required for the exit from latency (20, 33, 66). Finally, in contrast to observations from these primary neuronal cultures, infectious viruses can be detected in the eyes of HSV-1 latent rabbits and mice that have undergone reactivation by 24 hours, a timeframe that is inconsistent with Phase II of the model. Collectively, these data suggest that some neurons may be escaping latency very early following stress and that these events could lead to viral reactivation in a population of neurons. Our data show that this is a likely scenario, as we can detect VP16 protein expression in ∼15-20% of LUHMES cells stimulated to reactivate at 6 hpr. Nonetheless, our findings also likely highlight what researchers have long suspected in that HSV-1 reactivation is highly heterogeneous in nature and viral genomes harbored in the diverse populations of neurons perhaps utilize different de-repression mechanisms to achieve the uniform goal of producing progeny virus during a reactivation event.

Sawtell and Thompson have provided compelling evidence *in vivo* using heat stress-induced HSV-1 reactivation that viral replication is initiated by *de novo* synthesis of VP16 in a small subset of neurons (33, 66). Their model proposes that VP16 controls the exit from latency through mechanisms involving the de-repression of the VP16 promoter so that *de novo* synthesis of VP16 can activate viral IE gene expression. In this current work, we provide evidence that LTE is required for the de-repression of the VP16 promoter in a neuronal cell-specific manner. Considering the spatial distance between the VP16 promoter and the LAT region of the genome (over 20,000 base pairs) we suggest that CTCF insulators organize (some or all) viral genomes into a 3D chromatin structure that allows for the LTE to be proximal to the VP16 promoter. Our data support this hypothesis, as we have identified and functionally characterized a novel CTCF insulator (CTUL1) near the VP16 promoter locus that are enriched in both CTCF and cohesin proteins, suggesting the presence of 3D chromatin loops. Intriguingly, CTUL1, completely evicts both CTCF and cohesin by 2 hpr, and this eviction correlates to transient increased VP16 expression in a time-frame consistent with the exit from latency, suggesting that eviction of insulator proteins is required for the expression of VP16 and perhaps *de novo* synthesis of viral proteins. Importantly, this increase in VP16 occurs as early as 1 hpr and corresponds to the eviction of CTCF from insulators near VP16 and is independent of *de novo* viral proteins. Nonetheless, it is possible that the increased VP16 expression is not related to the eviction of CTCF, since the earliest timepoint that we measured CTCF eviction was at 2 hpr. However, previous work done by our lab has measured decreased CTCF enrichment on other CTCF insulators at 1 hpr, a time point not assessed in this study. Experimentally testing which happens first is difficult in both *in vivo* and *in vitro* models of reactivation used, since it requires either pharmacologically inhibiting reactivation, or using a recombinant virus that does not reactivate but maintains the 3D organization of the genome. For the former, we are currently exploring other *in vitro* methods of reactivation to help answer this question. For the latter, we provide compelling evidence that in the absence of the LTE, both VP16 expression remains unchanged following the application of a reactivation stimulus and that these viral genomes, while able to establish efficient latent infections, fail to reactivate and produce progeny viruses. Finally, our data also show that in the absence of the LTE, select genes including VP16 and ICP27 do increase in expression, while others do not, but these increases in expression are uncoupled to reactivation, suggesting that chromatin remodeling may still be occurring but that remodeling is not required for reactivation. This is an intriguing avenue for our future exploration because it outlines the heterogeneity of changes to HSV-1 genomes in response to stressors and further suggests that efficient reactivation may require multiple divergent pathways be activated.

Based on our results, we propose that the novel CTCF insulator CTUL1 acts to prevent the inappropriate activation of VP16 during latency by insulating the VP16 promoter from the LTE that could otherwise induce its expression (**Fig. 5**). We predict that these functional insulators are critical for maintaining the quiescent state of the viral genome within neurons. However, these insulators could alternatively facilitate the establishment of a repressive chromatin environment, characterized by the presence of heterochromatin markers such as H3K27me3. This is supported by the fact that cohesin can recruit and/or stabilize repressive histone-modifying complexes, such as those that methylate histone H3 on lysine 27 (H3K27me3). Upon reactivation, the chromatin landscape of the HSV-1 genome undergoes significant remodeling, and this could also lead to the de-repression of VP16. Nonetheless, the disruption of CTCF-mediated insulators (and cohesin) would be a key step in this process.

**Figure 5:**
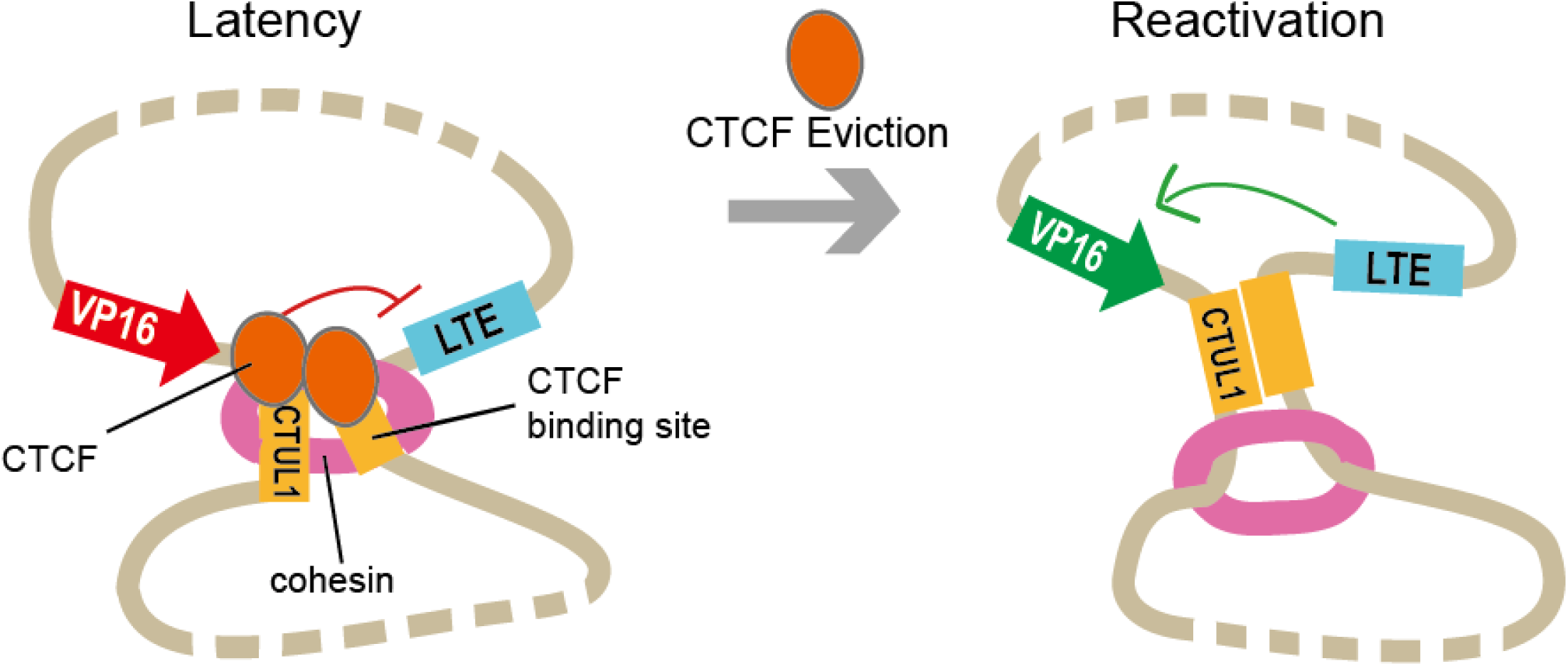
A model illustrating how CTCF insulators regulate VP16 *de novo* gene expression upon HSV-1 reactivation. Beige line indicates HSV-1 genome. Rectangles on the genome denote elements regulatory elements: blue: LAT enhancer; red/green: VP16 promoter; yellow: insulators. Red line denotes transcriptional repression while green line denotes activation. In latency, CTUL1 and another unidentified insulator are enriched with CTCF protein and anchored by the cohesin complex. This structure insulates LTE from turning on the VP16 promoter. In reactivation, CTCF proteins evict from the insulators and cohesin moves away. The loss of enhancer-blocking activity allows LTE to turn on the VP16 promoter.

While we are still exploring how CTCF is evicted, it is possible that reactivation signals, such as stress or immune suppression, may lead to the phosphorylation or proteolytic degradation of CTCF, weakening its binding to the viral genome. This could result in the loss of insulator function, allowing enhancer elements to interact with the VP16 promoter and initiate its transcription. The precise mechanisms by which CTCF function is modulated during reactivation remain an active area of research, but the evidence suggests that CTCF insulators play a dynamic role in the regulation of HSV-1 gene expression during the viral life cycle. Considering that the CTUL1 insulator near VP16 rapidly evicts CTCF, CTCF eviction is potentially a mechanism for de-repression of the VP16 promoter to assist the virus to exit latency.

## Methods

### Virus and cells

For mice Neuro-2a (N2a; ATCC CCL-131) and human epithelial kidney 293FT (HEK293FT; ThermoFisher R70007) cells used in transfection, cells were cultured in Dulbecco’s Modified Eagle’s Medium (DMEM; Corning 15013) with the presence of 10% fetal bovine serum (FBS; Corning 35011CV) and 1% antibiotic-antimycotic solution (Corning 30-004-CI). Cells were maintained in a humidified 37°C incubator supplemented with 5% CO_2_ and passaged regularly when the cells became nearly confluent. The undifferentiated Lund human mesencephalic (LUHMES; ATCC CRL-2927) cells were cultured in a cell cultural flask that was coated with 10 mg/ml poly-l-ornithine (Sigma P3655) and 1 mg/ml fibronectin (Corning 354008) before use. Cells were grown in 1:1 DMEM/F12 (Gibco 11330-032) with the presence of 1% N2 supplement (Gibco 17502048), 1% antibiotic-antimycotic and fibroblast growth factor (Peprotech 100-18B) at 37°C with 5% CO_2_. Cells were passaged when they reached 70% confluence. For herpes infection, the wild type HSV-1 strain, 17*Syn*+ (GenBank NC_001806), was a gift from J. Stevens. The recombinant 17ΔA strain, which has n.t.119,197-119,507 deletion from 17*Syn+,* was a gift from D. Bloom (University of Florida). These viruses were propagated in Vero cells (ATCC CCL-81) and titered in the Neumann lab.

### LUHMES infection

Undifferentiated LUHMES cells were seeded in a coated 6-well plate at 9 × 10^4^ cells per well in the growth medium as described above. One day later, differentiation was initiated by changing the medium to DMEM/F12, 1% N2 and 1% antibiotic-antimycotic supplemented with 10 mg/ml tetracycline hydrochloride (Sigma T7660), 100 μg/ml rhDGNF (R&D system 212GD) and 204 mM cyclic AMP (Sigma D0627). This differentiation medium was replaced again on day 1 and 3 post-differentiation. On day 5 post-differentiation, cells were infected by 17*Syn*+ or 17ΔA virus at a MOI indicated in each experiment in the presence of 50 μM Acyclovir (Sigma PHR1254) but absence of cAMP (infection medium), slowly rocked at 4°C for 1 hour, then moved to the 37°C incubator. 1 hour later, the medium was replaced with fresh infection medium. At 24 hours post infection, medium was again removed and replaced. At 48 hours post infection, medium was removed and changed to infection medium without Acyclovir. Medium was renewed every two days until cell harvest at 8 days post infection, when quiescence (latent-like infection) was established. For *in vitro* experiments involving reactivation, latent medium was replaced by fresh N2 medium with 10 mg/ml tetracycline and 1 μM wortmannin (Sigma W1628) (reactivation medium) for amount of time as indicated. For cycloheximide treatment during reactivation, latent LUHMES were pre-treated with cycloheximide (Sigma C7698) at a final concentration of 10 μg/mL in fresh latent medium for an hour. Then, medium was changed to reactivation media supplemented with 10 μg/mL cycloheximide for 6 hours of incubation.

### Quantitative reverse transcription-PCR

qRT-PCR was performed to compare the transcript level of VP16 during latency and reactivation. LUHMES cells were infected by 17*Syn+* or 17ΔA at MOI of 0.2 and maintained as described above in 6-well plates. Cells were harvested in 500 µL of TRIzol reagent (Ambion 15596026) per well at 8 days post-infection and 3, 6 and 24 hours post-reactivation. Total RNA was extracted according to the manufacturer’s protocol. qRT-PCR reactions were set up according to the manufacturer’s protocol. PCR for VP16 and GAPDH were performed with SuperScript III Platinum One-step qRT-PCR kit (Invitrogen 11732088) using the following PCR cycling conditions: 50 for 15 min, 95 °C for 5 min, and then 95 °C for 15 sec followed by 60°C for 30 sec (40×). VP16 transcripts were detected using Custom TaqMan™ Gene Expression Assay (AIQI074) and normalized to glyceraldehyde-3-phosphate dehydrogenase (GAPDH; ThermoFisher TaqMan gene expression assay Hs03929097_gl). PCR for ICP27, ICP8, gC and VP5 were performed with SuperScript™ III Platinum™ SYBR™ Green One-Step qRT-PCR Kit (Invitrogen 11736059) using the following PCR cycling conditions: 50 for 5 min, 95 °C for 5 min, followed by 40 cycles of 95°C for 15 sec and 60°C for 30 sec. Serial 10-fold dilutions of LUHMES or HSV-1 wt DNA were used to generate a standard curve for each plate, and the copy numbers of samples were determined based on this linear range.

### Viral genome copies and plaque assay

To determine relative genome quantity in lytic infection, Vero cells were infected by wt or 17ΔA strain at MOI of 0.1 in triplicate. At 2, 8, 18, and 24 hours post-infection, cells were harvested and total DNA were extracted by phenol-chloroform. To determine genome quantity in latent infection, LUHMES cells were quiescently infected by either wt or 17ΔA strain at MOI of 0.1 in triplicate. At 8 dpi (latency) or 48 hours post wortmannin addition (reactivation), cells were harvested and total DNA were extracted by phenol-chloroform, while the cell medium was saved at −80°C for plaque assay. Viral genome copy numbers were determined by qPCR using primers and probe specific for HSV-1 DNA polymerase (Table S2). qPCRs were performed using TaqMan™ Fast Universal PCR Master Mix (2×), no AmpErase™ UNG (Applied biosystems 4352042) on an Agilent AriaMx Real-Time PCR machine. The PCR cycling conditions were 95 °C for 10 min (1×) and then 95 °C for 15 sec, followed by 60°C for 1 min (40×). The copy numbers of DNA polymerase were then normalized to that of host GAPDH for a comparison between the two viruses. For plaque assay, Vero cells were seeded in a 24-well plate and maintained in 10% DMEM. When the cell confluency reaches 100%, medium was removed and cells were washed once by PBS (Corning 21-040-CM). Then, LUHMES cell medium taken at 8 dpi and 48 hpr were 1:10 diluted. 500 µL of undiluted and 1:10 diluted media were added to each well. The plate was incubated at 37°C with 5% CO_2_. After 2 days of incubation, medium was removed and 500 µL of 10% Neutral Buffered Formalin (epredia 5701) was added for cell fixation. After 30 min incubation at room temperature, NBF was removed and cells were stained by Gram Crystal Violet (Becton Dickinson 212525) overnight in dark. Plate was rinsed in water and the plaque-forming unit (PFU) was counted.

### Mouse ocular infections, genome copies and gene expression

Female Balb/c mice 4 to 6 weeks of age (Taconic) were anesthetized using an isoflurane-O_2_ chamber. Corneal scarification of each eye was performed in a 3×3 crosshatch utilizing a 25-gauge needle. Following scarification, 250,000 PFUs of wt or 17ΔA strain was applied to each eye. Mice were monitored daily through 28 dpi for signs of morbidity or mortality. Animals were considered latently infected at >28 dpi. For latent genome copies, TGs from 15 mice were harvested and total DNA was extracted by phenol-chloroform. Viral genome copies number were determined by qPCR targeting HSV-1 DNA polymerase and then normalized to copies of GAPDH (ThermoFisher TaqMan gene expression assay Mm99999915_g1). For gene expression study, viral reactivation was induced by the explant method as previously described [43]. In brief, mice TGs were harvested from >28 dpi mice, placed in DMEM supplemented with 1% FBS and 1% antibiotic-antimycotic solution for 2, 6 or 48 hours. Then, total RNA was extracted in TRIzol reagent according to the manufacturer’s protocol. qRT-PCR was performed as described above to determine RNA transcript level.

### Immunofluorescence Microscopy

LUHMES cells were grown to ∼50% confluency and differentiated for five days on 4-well chamber slides (Lab-Tek). Fully differentiated LUHMES cells were infected at an MOI of 1 in the presence of 50 μM Acyclovir with either wt or 17ΔA viruses. Latent infection was established as described in the “LUHMES infection” section. On day 9 post-infection, viruses were reactivated by 1 μM wortmannin. At 6 hours post-reactivation, cells were fixed with 4% PFA (Alfa Aesar J61899) for 10 min. Cells were washed with PBS and then incubated at room temperature for 5 min in 0.25% Triton X-100 in PBS. Blocking was done with 1% BSA in PBS for 1 hour at 37°C. VP16, mouse monoclonal primary antibody (1:1000; Abcam 110226) incubations were performed for 1 hour at room temperature in 1% BSA in PBS, followed by 5 washes with PBS for 10 min each. Secondary antibody anti-rabbit Alexa Fluor 488 (Invitrogen A21206) incubations were performed for 30 min at room temperature, followed by 5 washes in PBS for 10 min each. Slides were mounted with Prolong Gold with DAPI (Life Technologies). Images were captured with a Zeiss Axiomager Z2 upright fluorescent microscope using ZenPro software.

### CTCF insulator prediction

CTCFBSDS 2.0, a database of experimentally determined CTCF binding sites across several species was developed to predict CTCF binding sites *in silico*. Core CTCF binding motifs were represented by position weight matrices (PWM), and this database uses six PMW to search for CTCF motifs in any given sequence. CTCF binding at 2 kb region upstream of the VP16 gene (n.t. 101,200-103,600) in 17*Syn*+ genome was examined for putative CTCF binding sites. Three fragments of size around 800 bp were input to the prediction tool, and only output with a PWM score > 2.5 were considered as significant hits. PCR primers flanking these putative CTCF binding sites were designed and used for subsequent cloning. Each primer includes three parts: a 6-bp overhang sequence, a restriction site for cloning, and a 14-20 bp region of 17*Syn*+ genome.

### Plasmid construction

Plasmid encompassing the enhancer in the LAT region (LTE) and each of the putative CTCFBS were constructed to test the enhancer-blocking activity of these putative insulators. To begin, LTE (n.t. 118,888-119,477) was PCR amplified and inserted in between the KpnI and SacI restriction enzyme digestion sites in pGL3-control vector (Promega E1741), a reporter vector with a firefly luciferase gene under the control of a SV40 promoter. Then, each of the putative CTCFBS fragment was amplified from the 17*Syn*+ genome using primers described in Table S3 and inserted between LTE and the SV40 promoter. The sequences of all plasmids generated were confirmed by Sanger sequencing (Functional Biosciences). Plasmid encompassing LTE and VP16/VP5 promoter were constructed in the pGL3-basic vector that lacks any promoter or enhancer sequence (Promega E1751). LTE was PCR amplified and inserted in between the KpnI and SacI restriction enzyme digestion sites. Then, VP5 or VP16 promoter sequences were PCR amplified with the primers described in Table S3 and inserted between LTE and the luciferase gene. For the LTE-CTUL1-VP16/VP5 promoter construct, CTUL1 was amplified using primers described in Table S3 and inserted between LTE and the VP16/VP5 promoter. The sequences of all plasmids generated were confirmed by Sanger sequencing (Functional Biosciences).

### Transfection and Luciferase assay

The transient transfection protocol was adapted from the Invitrogen *Lipofectamine ™ 3000 Reagent Protocol.* Briefly, HEK293FT or N2a cells were seeded in a 24-well plate at a density of 8× 10^5^ cells per well in 10% DMEM 2 days before transfection. At the time of transfection, the cells were 70-80% confluent and medium were replaced by 1% DMEM. 500 ng of experimental plasmid purified by maxiprep (Promega A2393) and 5 ng of pMCS-Green Renilla Luc vector (ThermoFisher 16152) was co-transfected in each transfection reaction, in the presence of 0.75 μL of Lipofectamine ™ 3000 Reagent and 1 μL of P3000 Reagent (Invitrogen L3000008), according to the manufacturer’s protocol. The plasmid-Lipofectamine ™ 3000 complex was added directly on top of the cells and incubated for 48 hours at 37°C with 5% CO_2_. To visualize the transfection efficiency, 500 ng of plasmid encoding GFP was included as one of the experimental plasmids. Transfection reaction was performed in technical triplicate for each experimental plasmid. After 48 hours of incubation, cells were harvested and washed in PBS for luciferase assay. The luciferase assay was performed using a Dual-Luciferase® Reporter 1000 Assay System (Promega E1980). Washed cells were lysed in Passive Lysis Buffer, frozen at −80°C for 1 hour, and thawed. Each transfection reaction was mixed with 100 μL of LARII and 100 μL of Stop&Glo® reagent, and luciferase levels were determined by a Tecan SPARK Microplate Reader and SparkControl Software.

### ChIP-qPCR

Latently infected LUHMES were harvested at 8 dpi and washed in PBS with protease inhibitors (Aprotinin, Leupeptin and PMSF). Cells were crosslinked by formaldehyde (RICCA RSF0010-500A) at 1% final concentration and quenched by glycine (Fisher chemical G46-1). Samples were washed with PBS three times and resuspended in ChIP SDS-lysis buffer with protease inhibitors. After 30 min incubation on ice, samples were sonicated to shear the chromatin into 300-800 bp size fragments, verified by agarose gel electrophoresis. The chromatin was pre-cleared by adding Protein A agarose/salmon sperm DNA-50% slurry (EMD Millipore 16-157) and rocked at 4°C for 2 hours. Samples were centrifuged to sediment slurry and the supernatant was split to four equal aliquots: the first saved as the “input”, the second incubated with 5 μL of CTCF (EMD Millipore 07-729), the third with 15 μL of STAG2 (Cell signaling 5882), and the last with 2 μL of negative control antibody IgG (EMD Millipore PP64B) overnight at 4°C. To capture the antibodies, Protein A agarose/salmon sperm DNA was added and incubated for 2 hours at 4°C. The antibody-chromatin complexes were washed in 150 mM low salt buffer, 500 mM high salt buffer, 0.25 M LiCl buffer and TE buffers sequentially, and eluted as the “bound” fraction. The “input” and “bound” fractions were treated with 5 M NaCl to reverse crosslinks, followed by RNase A (Thermo Scientific EN0531) and Proteinase K (Thermo Scientific EO0491) incubation. DNA was purified by column purification (Qiagen 28106). qPCR was performed as described above.

### CUT & RUN-qPCR

CUT&RUN in latently infected LUHMES was performed with the CUT& RUN Assay Kit (Cell signaling 86652) following manufacturer’s protocol. Briefly, 8 days post-latent infection, cells in each well were harvested and washed in 1×Wash Buffer separately. One third of a well was saved as the “input” fraction at 4°C. The remaining two third of the cells were mixed with activated Concanavalin A magnetic beads and split in half. One half was added with 5 μL of CTCF antibody (Cell signaling 3418s) and the other with 2 μL of negative control antibody IgG (Cell signaling 66362). After overnight incubation at 4°C, cell:bead suspension was washed twice with Digitonin Buffer, resuspended in pAG-MNase pre-mix, followed by 1 hour rocking at 4°C. The cell:bead suspension was washed twice in Digitonin Buffer, and the pAG-MNase was activated to digest by cold 100 mM Calcium Chloride at 4°C for 30 min. Then, 1×Stop Buffer was added to stop the digestion and release DNA fragments. Beads were discarded and the supernatants were saved as the “Bound” or “IgG” chromatin fraction. To prepare for the qPCR analysis, the input fraction was first sonicated to 100-300 bp size fragments. Then, all three fractions were purified using the DNA Purification Buffers and Spin Columns (Cell signaling 14209) and proceeded to qPCR. To determine whether the CTUL1 was enriched with CTCF, primers and probe targeting the CTCF binding site were designed. qPCRs were performed using TaqMan™ Fast Universal PCR Master Mix (2×), no AmpErase™ UNG (Applied biosystems 4352042) on an Agilent AriaMx Real-Time PCR machine. The PCR cycling conditions were 95 °C for 10 min (1×) and then 95 °C for 15 sec followed by 60 °C for 1 min (40×). Serial 10-fold dilutions of LUHMES or HSV-1 wt DNA were used to generate a standard curve for each plate, and the copy numbers of samples were determined based on this linear range.

### ChIP and CUT&RUN validation

All CTCF and STAG2 binding in CUT & RUN and ChIP-qPCR were validated by determining the Bound/Input ratio of a cellular positive binding control and compared that to IgG. Primer and probe sequences are listed in Table S2. Only samples that have more than 2-fold abundance of CTCF or STAG2 compared to IgG bound at the positive control site were used in further viral targets analyses.

### Statistics analysis

All bar graphs are presented as the mean with standard deviation, unless otherwise noted in the figure legends. For luciferase assays, ChIP-qPCR and gene expression assay, we applied two-tail distributed Student’s *t*-tests with two-sample unequal variance. As shown in the figures, * denotes *p* < 0.05, ** *p* < 0.005, *** *p* < 0.0005, ns denote not significant or *p*> 0.05. For CUT&RUN-qPCR and mice studies, one-way analysis of variance (ANOVA) was performed on fold changes. * denotes *p* < 0.05, ** *p* < 0.005, *** *p* < 0.0005, ns denote not significant or *p*> 0.05.

## Acknowledgements

This work was supported by the grants NIH/NIAID R01AI134807 (DMN), NIH/NEI 2T32EY027721 (MAS), the Core Grant for Vision Research from the NIH to the University of Wisconsin-Madison (P30 EY016665), the McPherson Eye Research Institute Grant Summit Program and an unrestricted grant from Research to Prevent Blindness (Department of Ophthalmology, University of Wisconsin). The funders had no role in study design, data collection and analysis, decision to publish, or preparation of the manuscript.

