## supplemental figure 1 for "In HSV-1, the LAT Enhancer Drives Pre-IE VP16 Transcription to Initiate Reactivation"

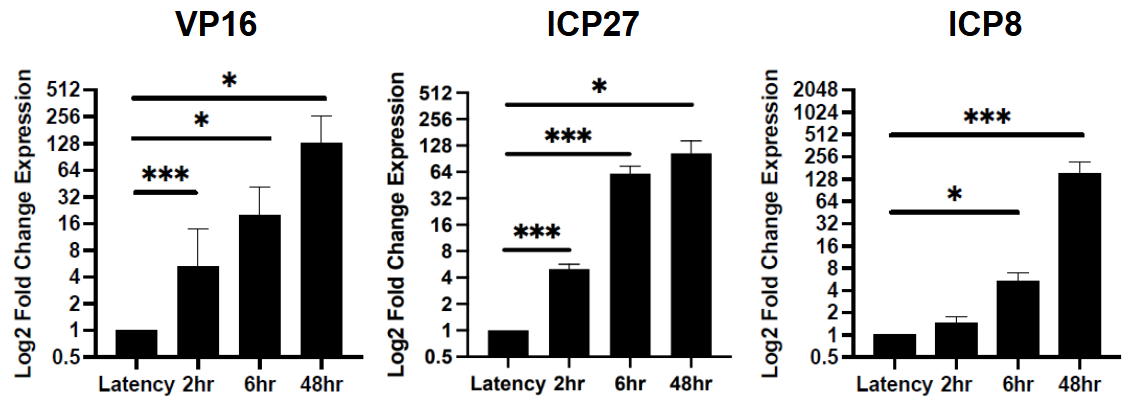

**Supplemental Figure 1**: wt gene expressions upon reactivation were quantified by RT-qPCR in latently infected mice TG. After latency was established, reactivation was induced by explant of TG. Total RNA was extracted at 2, 6 and 48 hours post-explant. Relative values for each gene were normalized to host GAPDH expression and were plotted as fold change in expression relative to latency (set to 1). n=6 and each n represents one TG. **p*<0.05, ** *p*<0.005, ****p*<0.0005 following one-way ANOVA.
