## supplemental figure 2 for "In HSV-1, the LAT Enhancer Drives Pre-IE VP16 Transcription to Initiate Reactivation"

A

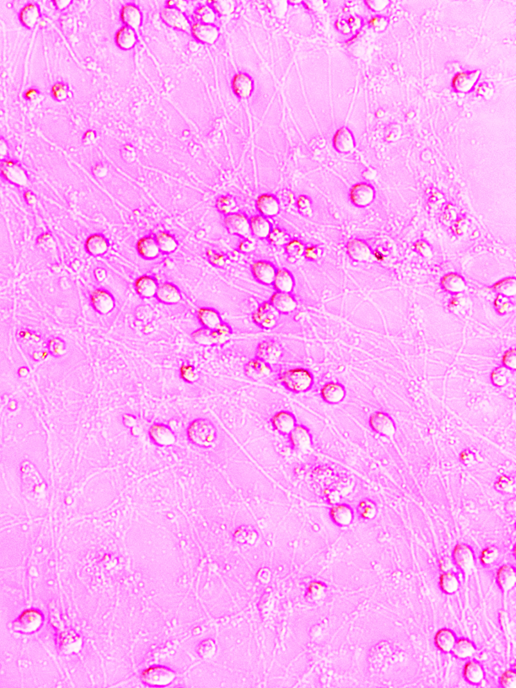

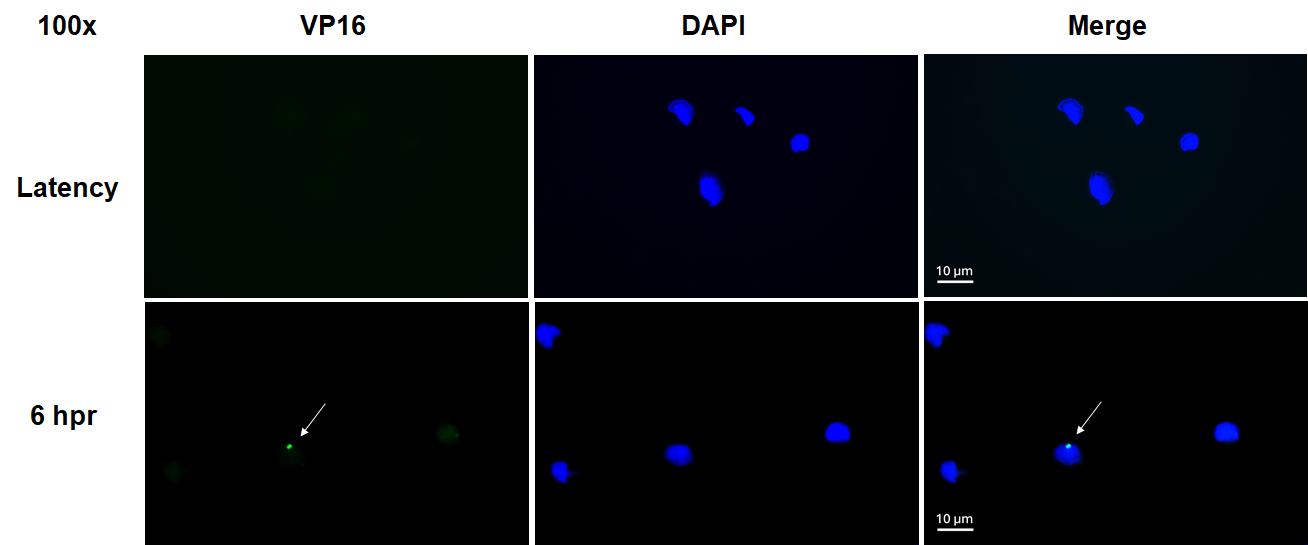

B

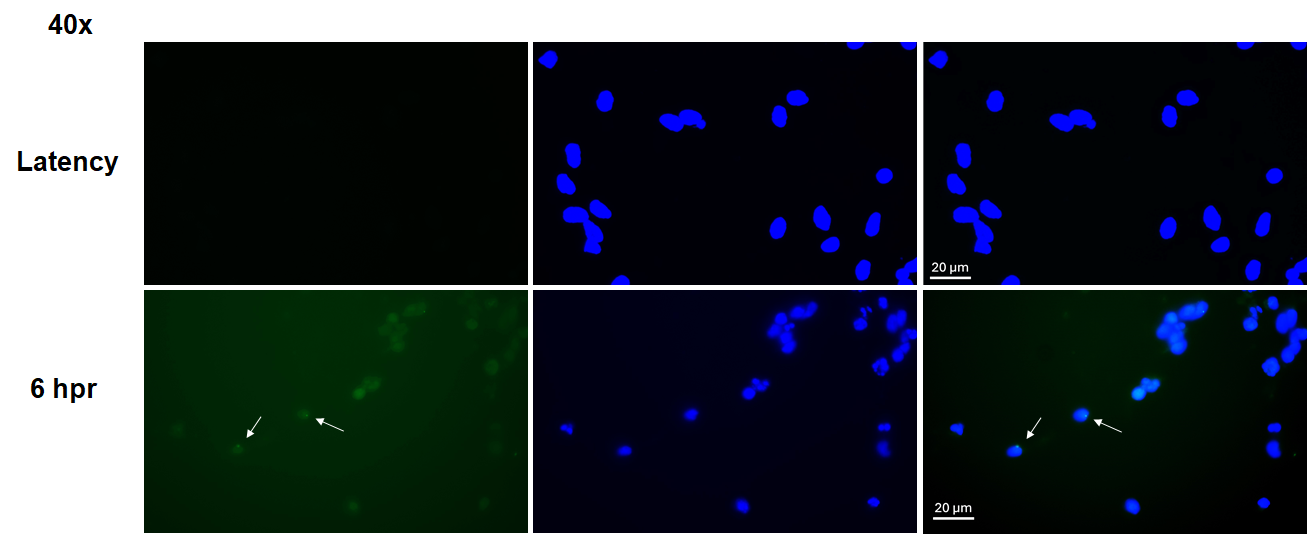

**Supplemental Figure 2**: **A**. Phase-contrast microscopy image of latently infected LUHMES induced to reactivate with wortmannin. Representative image taken at 9 dpi at 200x magnification to show axon growth and retention. **B**. Representative images showing immunofluorescence of the VP16 protein in LUHMES at 0 (latency) and 6 hours post-reactivation. Green: VP16 protein; Blue: DAPI staining of cell nucleus. Upper panel: images taken at 100x magnification; lower panel: 40x magnification. Scale bar represents 10 µm on 100x images and 20 µm on 40x images. Arrows pointing at VP16 foci.
