## supplemental figure 3 for "In HSV-1, the LAT Enhancer Drives Pre-IE VP16 Transcription to Initiate Reactivation"

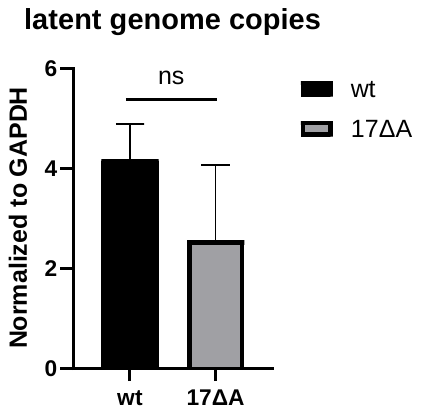

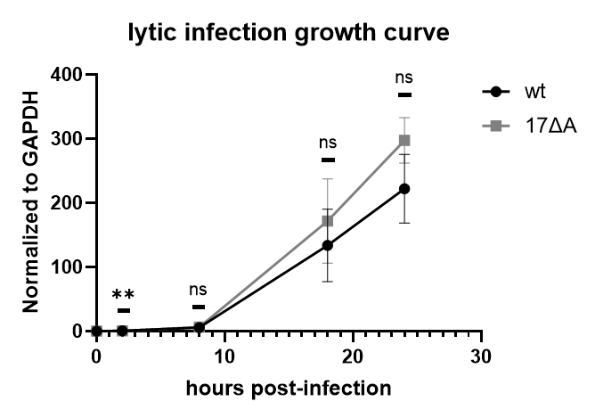
Supplemental Figure 3:

A

B

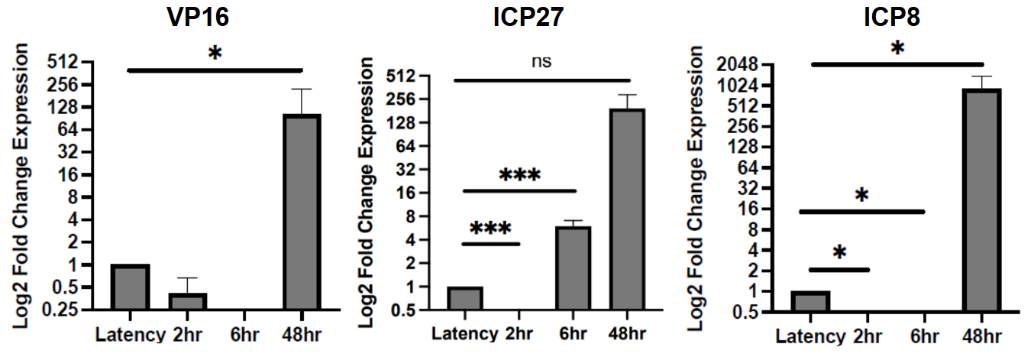

C

**Supplemental Figure 3**: 17∆A showed the same trend of gene expression in early reactivation *in vivo* as *in vitro*. **A**. Relative quantity of wt and 17∆A HSV-1 genome copies was determined during lytic infection in Vero cells. Cells were harvested at 2, 8, 18 and 24 hours post-infection without Acyclovir. Relative genome load was determined by qPCR copy number of DNA polymerase normalized to that of host GAPDH. n=3 for each virus at each time point. Statistical significance between wt and 17∆A was calculated by Student’s t-tests. **B.** wt and 17∆A had comparable genome copies in latent mice TG (>28 dpi) determined by qPCR of DNA polymerase/GAPDH. Plot displays mean with SEM. n=15 and each n represents TG pooled from one mouse. Statistical significance was calculated by one-way ANOVA. **C.** 17∆A gene expressions were quantified by qRT-PCR in mice TG. After latency was established, reactivation was induced by explant of TG. Total RNA was extracted at 2, 6 and 48 h post-explant. Relative values for each gene were normalized to host GAPDH expression and were plotted as fold change in expression relative to latency (set to 1). n=6 and each n represents one TG. **p*<0.05, ** *p*<0.005, ****p*<0.0005 following one-way ANOVA.
