## supplemental figure 4 for "In HSV-1, the LAT Enhancer Drives Pre-IE VP16 Transcription to Initiate Reactivation"

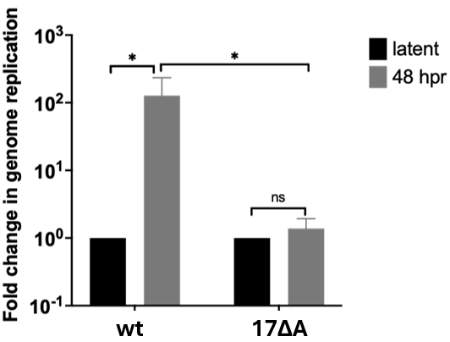
Supplemental Figure 4:

A

B

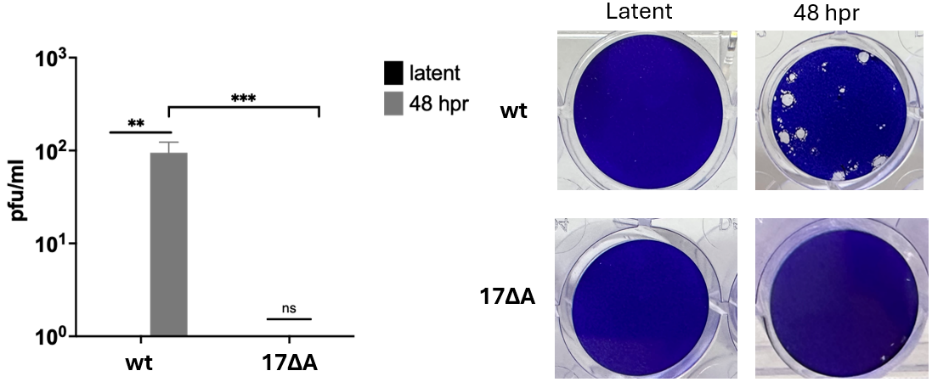

**Supplemental Figure 4**: 17∆A fails to reactivate. **A.** 17∆A HSV-1 had significantly fewer genome copies compared to wt at 48 hour post-reactivation. Relative genome load was determined by qPCR copy number of DNA polymerase normalized to that of host GAPDH. n=3 for each virus at each time point. Statistical significance was calculated by Student’s t-tests. **B.** Representative images of plaque assays performed using 1:10 diluted supernatant taken from LUHMES infected by wt and 17∆A. n=6 for each virus. ***p*<0.005, ****p*<0.0005 by Student’s t-tests.
