## supplemental figure 5 for "In HSV-1, the LAT Enhancer Drives Pre-IE VP16 Transcription to Initiate Reactivation"

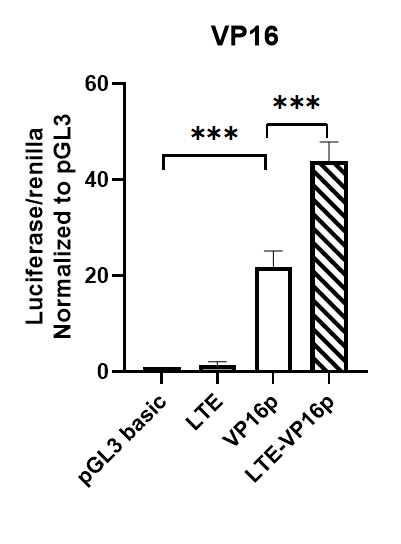

**Supplemental Figure 5:** LTE is not required for the activation of the VP16 promoter in HEK293FT cells. All transfections were completed in triplicate wells and were repeated three times biologically. All luciferase values were normalized to the pGL3-basic vector (set to 1). *p* values were determined by unpaired two-tailed Student’s t-tests with unequal variances. *** *p<*0.0005.
