## supplemental figure 6 for "In HSV-1, the LAT Enhancer Drives Pre-IE VP16 Transcription to Initiate Reactivation"

A

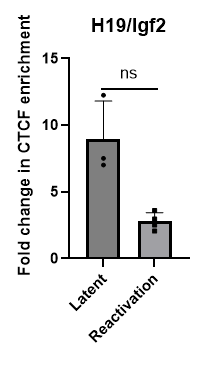

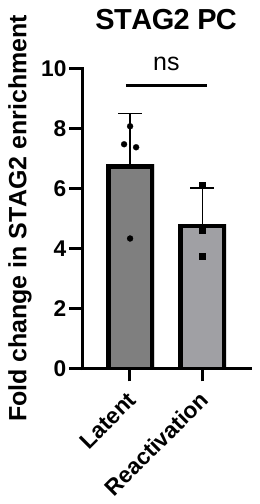

B

**Supplemental Figure 6**: There was no significant change in CTCF and cohesin enrichment at host positive control region between latency and 2hpr. **A.** ChIP-qPCR for CTCF was done as reported at host H19/Igf2. **B.** ChIP-qPCR for STAG2 at the host Positive Control region on chr.1. Values are plotted as fold enrichments over IgG in latency vs. 2 hpr. Difference between latency and reactivation was determined statically by unpaired two-tailed Student’s t-test.
