## Supplemental Table 1 for "In HSV-1, the LAT Enhancer Drives Pre-IE VP16 Transcription to Initiate Reactivation"

| **Site PWM** | **Site Sequence** | **Location** | **Site orientation (+/- strand)** | **Score** | **Site name** |
| --- | --- | --- | --- | --- | --- |
| REN_20 | CTCGCCAACAGCGGGCTCAA | 101,634-653 | - | 8.14 | **A** |
| EMBL_M1 | AGCCCGCTGTTGGC | 101,637-650 | + | 2.68 |  |
| EMBL_M2 | GGTGGTGCA | 101,767-775 | + | 6.98 | **B** |
| EMBL_M2 | AGTACAGCA | 102,013-021 | + | 10.86 | **C** |
| MIT_LM7 | TAGCCAGGAGACGGGACCGC | 103,454-473 | - | 4.92 | **D** |

**Supplemental Table 1:** 2 kb region upstream of the VP16 gene (n.t. 101,200-103,600) was input to CTCFBSDS 2.0, a web tool to search for CTCF-binding sites core motifs [48, 49]. Motifs are represented by position weight matrices (PWM).  Nucleotide locations, strand orientation and scores are presented.  For simplicity, each putative binding site name was designated A-D. Only motifs with PWM score >2.5 were listed in the table and shown on the genome.
