## Supplemental Table 2 for "In HSV-1, the LAT Enhancer Drives Pre-IE VP16 Transcription to Initiate Reactivation"

**Supplemental Table 2**: Primer sequences used in qPCR and reverse transcriptase-qPCR.

| **Experiment** | **Target** | **Primer DNA sequences (5’ to 3’)** | **Location in genome** |
| --- | --- | --- | --- |
| ChIP and CUT&RUN-qPCR | CTUL1 | For: TAACACACACGCCGGCAA  Rev: CGGTTGTCGTCTGATTATTTGGT  Probe: 5'FAM-CTGGCTAGTTCTTTCCCCCA-3’BHQ | 103,412-103,511 |
|  | CTRL2 | For: CCGCGGCTCTGTGGTTA  Rev: GGATGCGTGGGAGTGGG  Probe: 5'FAM- ACACCAGAGCCTGCCCAACA  TGGCA- 3’BHQ | 120,455-120,643 |
|  | human H19/Igf2 | For: CTCACACATCACAGCCCAAG  Rev: ATGAGCGTCCTATTCCCAGA | (NCBI accession no. NG_041945.2) 3,990-4,105 |
|  | STAG2 positive Control | For: AGACCACGACGATTGGAGAC  Rev: CCCTGGACACTGGCATCTAC | human chr.1: 153483538-153483657 |
|  | gC | For: CCTTGCCGTGGTCCTGTGGA  Rev: GGTGGTGTTGTTCTTGGGTTTG | 96,332-96,644 |
| Gene expression RT-qPCR | Human GAPDH | ThermoFisher TaqMan gene expression assay Hs03929097_gl | human chr.12: 6534405 – 6538375 on Build GRCh38 |
|  | VP16 | ThermoFisher TaqMan gene expression assay AIQ1074 | Custom design |
|  | ICP27 | For: GCCCGTCTCGTCCAGAAG  Rev: GCGCTGGTTGAGGATCGTT | 113,945-114,035 |
|  | ICP8 | For: CTCAAAGCCGCTCTCCAC  Rev: ATGGAGACAAAGCCCAAGAC | 61,869-62,054 |
|  | gC | For: CCTTGCCGTGGTCCTGTGGA  Rev: GGTGGTGTTGTTCTTGGGTTTG | 96,332-96,644 |
|  | VP5 | For: AGCTCCGGAAACTTGGTACA  Rev: CTCCTTAGCACGATCGAGGT | 40,270-40,463 |
|  | Mouse GAPDH | ThermoFisher TaqMan gene expression assay Mm99999915_g1 | mouse chr.6: 125161338 - 125166511 on Build GRCm38 |
| Genome copies qPCR | DNA polymerase | For: AGAGGGACATCCAGGACTTTGT  Rev: CAGGCGCTTGTTGGTGTAC  Probe: 5’FAM-ACCGCCGAACTGAGCA-3’BHQ | 65,880-65,953 |
