## Supplemental Table 3 for "In HSV-1, the LAT Enhancer Drives Pre-IE VP16 Transcription to Initiate Reactivation"

**Supplemental Table 3**: Primer sequences used in the generation of reporter constructs.

| **Name** | **PCR primer sequence**  **(lowercase indicates digestion site)** | **Restriction digestion site** | **Amplicon genomic location** |
| --- | --- | --- | --- |
| LTE | For: CTCTCT ggtacc GTCGGCGACATCCTCCCCCT  Rev: CCTTCT gagctc GGTGTCTAACCTACCTGGAAACGCGG | KpnI and SacI | 118,888-119,477 |
| Site A | For: CTCTCT gctagc GTGAATCCCCCGTCAAAC  Rev: CCTCCT agatct CCGCCTTTGCGGACGTG | NheI and BglII | 101,418-101,766 |
| Site B | For: CTCTCT agatct CAGCCGGCGCGCTG  Rev: CCTCCT ctcgag AGACGGTCCAGTGGCTCTC | BglII and XhoI | 101,654-101,826 |
| Site C | For: CTCTCT gctagc GGTGGCCAGGTACGTCAT  Rev: CCTCCT agatct ATCAACGGGTGGGTGTGC | NheI and BglII | 101,933-102,232 |
| Site D (CTUL1) | For: CTCTCT gctagc TGGTTTCGAATCTACACGACA  Rev: CTCTCT agatct CACCGGTTCGTATCCACAAT | NheI and BglII | 103,375-103,560 |
| CTRL2 positive control | For: CTCTCT gctagc GTGCCTTTGCACACCAAC  Rev: CTCCTCCT agatct ACCACAGACGTGGGTGTTG | NheI and BglII | 120,000-120,200 |
| VP5 promoter | For: CTCTCTC gctagc GACCAGGGCCATCTTGAATG  Rev: CCTCCT ctcgag TCCACCGAAAGACACGACG | NheI and XhoI | 40,631-40,857 |
| VP16 promoter | For: CTCTCTC gctagc GTTCATTCGGTGTTGGCGTT  Rev: CCTCCT ctcgag GATATCGGGCTTTTGATGCG | NheI and XhoI | 105,106-105,450 |
| Random sequence near CTUL1 | For: CTCTCTC gctagc GGGACCCCAGCGAGCT  Rev: CCTCCT ctcgag ATTCTTGACCTGACGGATCG | NheI and XhoI | 102,022-102,316 |
